# An isogenic collection of pluripotent stem cell lines with elevated α-synuclein expression

**DOI:** 10.1101/2022.02.10.479903

**Authors:** Ammar Natalwala, Ranya Behbehani, Ratsuda Yapom, Tilo Kunath

**Affiliations:** UCL Queen Square Institute of Neurology, Department of Neuromuscular Diseases, Queen Square House, London, WC1N3BG; Victor Horsley Department of Neurosurgery, National Hospital for Neurology & Neurosurgery Queen Square, London, WC1N 3BG; Centre for Regenerative Medicine, Institute for Regeneration and Repair, School of Biological Sciences, The University of Edinburgh, Edinburgh BioQuarter, 5 Little France Drive, Edinburgh, EH16 4UU

**Author notes:** **Corresponding authors** Mr Ammar Natalwala *BSc(Hons), MBChB, MRCS (Eng), PhD*, NIHR Clinical Lecturer in Neurosurgery, Tilo Kunath *PhD. **Funding** AN was funded by the Wellcome Trust Research Training Fellowship (203646/Z/16/Z). TK was funded by an MRC grant (MR/J012831/1).

**Keywords:** Human pluripotent stem cells, α-synuclein, synucleinopathy, isogenic cell lines, cellular differentiation, neurogenesis, transcriptomics, Parkinson’s disease, dementia with Lewy bodies

## Abstract

α-Synuclein (αSyn) is a small, disordered protein that becomes aggregated in Lewy body diseases, such as Parkinson’s disease (PD) and dementia with Lewy bodies (DLB). Human induced pluripotent stem cells (hiPSCs) potentially provide a tractable disease model to monitor early molecular changes associated with PD/DLB. We and others have previously derived hiPSC lines from patients with duplication and triplication of the *SNCA* gene, encoding for αSyn. It is now recognised that to perform meaningful disease modelling with these hiPSC lines, it is critical to generate isogenic control cell lines that lack the disease causing mutations. In order to complement the existing and emerging hiPSC models for PD/DLB, we have generated an allelic series of αSyn over-expressing hESC lines on the same isogenic background. An unresolved question is whether pluripotent stem cell lines, with elevated levels of αSyn, can undergo efficient differentiation into dopaminergic and cortical neurons to model PD and DLB, respectively. We took advantage of our isogenic collection of hESC lines to determine if increased expression of αSyn affects neural induction and neuronal differentiation. Clonal hESC lines with significantly different levels of αSyn expression proliferated normally and maintained expression of pluripotent markers, such as OCT4. All cell lines efficiently produced PAX6^+^ neuroectoderm and there was no correlation between αSyn expression and neural induction efficiency. Finally, global transcriptomic analysis of cortical differentiation of hESC lines with low or high levels of αSyn expression demonstrated robust and similar induction of cortical neuronal expression profiles. Gene expression differences observed were unrelated to neural induction and neuronal differentiation. We conclude that elevated expression of αSyn in human pluripotent stem cells does not adversely affect their neuronal differentiation potential and that collections of isogenic cell lines with differing levels of αSyn expression are valid and suitable models to investigate synucleinopathies.

## Introduction

Multiple lines of evidence have implicated αSyn as a major pathological driver in PD (Chartier-Harlin et al., 2004; Polymeropoulos et al., 1997; Singleton et al., 2003; Spillantini et al., 1997). The genetic forms of PD have led to the development of patient-derived and engineered pluripotent stem cell-derived models to gain mechanistic insights into synucleinopathies (Singh Dolt et al., 2017). A key question for the validity of such models is whether mutant or elevated αSyn expression disrupts early neural induction or neuronal differentiation, thereby limiting later phenotypic analysis and disease modelling. Current data in the literature is conflicting, with some studies proposing that high levels of αSyn alter cell fate and differentiation (Oliveira et al., 2015; Schneider et al., 2007), whereas others have found no evidence for impaired neurogenesis (Brazdis et al., 2020; Devine et al., 2011; Prots et al., 2018). In this study, we aimed to generate an isogenic collection of hESC lines with differing levels of αSyn expression, and use this to systematically investigate if elevated αSyn expression affects cortical neuron differentiation. If either the efficiency of neural induction or fidelity of neuronal differentiation is affected, then this may be a source of bias in studies comparing cultured neurons with differential αSyn expression. Isogenic cellular models provide the best experimental system to test this directly.

Studies utilising αSyn null mouse models showed no major differences in neuronal development and overall brain structure (Abeliovich et al., 2000; Greten-Harrison et al., 2010; Specht and Schoepfer, 2001). However, subtle differences in dopaminergic neurotransmission were reported and these were age-dependent (Al-Wandi et al., 2010; Anwar et al., 2011; Connor-Robson et al., 2016). Rodent models employing transgenic over-expression of wild-type αSyn have reported impaired adult neurogenesis, but have not determined whether this is a neurodevelopmental defect from birth or a toxic effect of αSyn over-expression (Winner et al., 2012). We have previously shown that hiPSCs with an *SNCA* triplication mutation have a two-fold increase in αSyn protein levels, and this did not significantly impair dopaminergic neuronal differentiation (Devine et al., 2011). This was also shown by other independent groups using hiPSCs harbouring the *SNCA* triplication (Byers et al., 2011; Lin et al., 2016) and *SNCA* duplication mutations (Brazdis et al., 2020; Prots et al., 2018). However, other studies using *SNCA* triplication mutation-derived hiPSCs reported reduced neuronal differentiation capacity and impaired neurite outgrowth (Flierl et al., 2014; Oliveira et al., 2015). Furthermore, other studies using lentiviral-mediated Dox-inducible expression of αSyn, in human iPSC-derived neuronal progenitors, reported altered cell fate and impaired differentiation of neural stem cells into neurons (Schneider et al., 2007; Zasso et al., 2018). The cellular and rodent data described so far suggests conflicting roles for αSyn in neurogenesis, highlighting the need for work to clarify this point.

In this study, we generated an allelic series of clonal isogenic hESC lines expressing a broad range of αSyn, including lines expressing the protein at supraphysiological levels. We then set out to determine whether elevated αSyn expression impairs cortical neuron differentiation of human pluripotent stem cells. We show, using marker analysis during multiple rounds of differentiation and unbiased transcriptomic analysis, that cortical neuron differentiation is not impaired by increased expression of αSyn. The strength of our approach is the use of an isogenic collection of hESC lines with differing and sustained αSyn expression during differentiation, robust cortical neuron differentiation protocols and unbiased analysis of the transcriptome before and after differentiation. This study provides a valuable collection of cell lines for the neuroscience community and provides important evidence for the validity of hESC/iPSC disease models with elevated levels of αSyn expression.

## Results

### Generation of clonal isogenic hESC lines with differing levels of αSyn expression

An allelic series of clonal transgenic hESC lines were established using a human *SNCA* cDNA expression cassette driven by the pCAG promoter (Figure 1A), reported to maintain stable and ubiquitous transgene expression across diverse cell types (Hitoshi et al., 1991; Liew et al., 2007). Multiple puromycin-resistant clones were established on a parental Shef4 hESC line and examined for *SNCA* expression using qRT-PCR (Figure 1B). In an undifferentiated state, 10 clones, including S9 and S34, had similar *SNCA* expression to Shef4, whereas 19 clones, including S8 and S37 had elevated expression of *SNCA* (2-fold to 30-fold) relative to the parental cell line (Figure 1B). V2 and V39 control lines, transfected with a control plasmid lacking *SNCA*, expressed similar levels of *SNCA* to the parental cell line, Shef4. Clones were selected based on *SNCA* mRNA expression and placed into low (Shef4, S9, S34) and high (S8, S37) groups. Each of these lines similarly expressed the pluripotency marker, OCT4, and maintained hESC morphology in their undifferentiated state (Figure 1C). Elevated expression of αSyn protein was confirmed for selected clones using immunohistochemistry and western blotting (Figure 1C,D).

**Figure 1.**
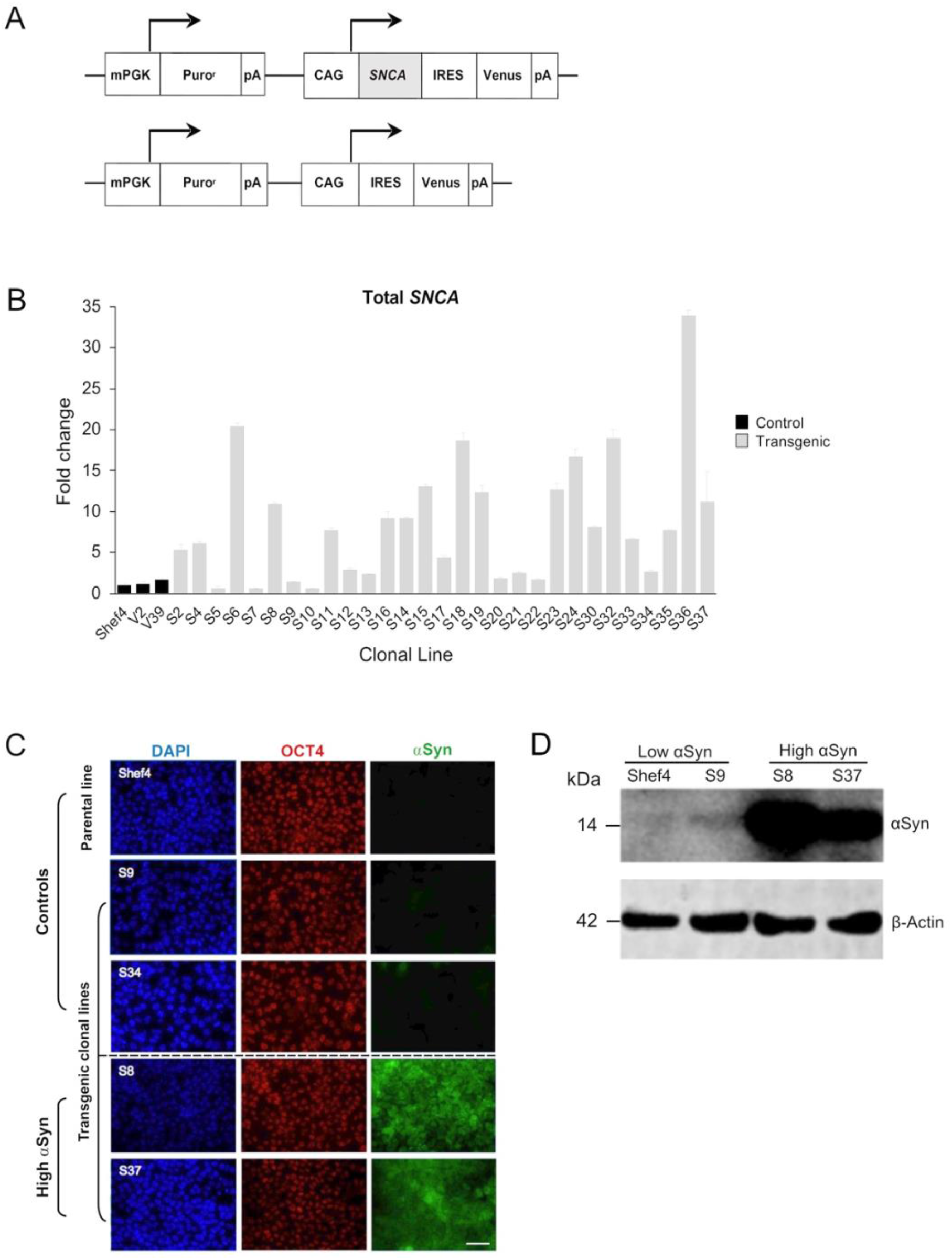
Establishment of clonal hESC lines over-expressing αSyn. (A) Schematic of pCAG-SNCA-IRES-Venus and pCAG-IRES-Venus constructs transfected into Shef4 hESCs to generate clonal lines. (B) Quantitative RT-PCR measuring total *SNCA* expression (mRNA) levels in multiple self-renewing undifferentiated transgenic clones generated from parental Shef4 hESC lines. Data was normalised to 18S rRNA levels, and shown relative to expression in the parental hESC Shef4 line. Each bar represents the mean and standard deviation of 3 technical replicates. (C) Representative immunocytochemistry images of undifferentiated hESC clones derived from the parental cell line Shef4, and transgenic clonal lines S9, S34, S8, and S37, stained for DAPI (blue), OCT4 (red), and αSyn (green). Scale bar = 50 μm. (D) Western blot for total αSyn and β-actin in undifferentiated clonal lines for Shef4 and S9 (low αSyn), and S8 and S37 (high αSyn) clones.

The impact of αSyn on cell proliferation was assessed using an MTS assay. Over a 10-day period, six low αSyn lines (S9, S12, S13, S17, S22, S34) and 3 high αSyn lines (S8, S36, S37) were examined and each showed a similar rate of proliferation (Supplementary Figure 1). There was no significant difference in the proliferation rate of low and high αSyn groups and linear regression analysis showed no correlation between *SNCA* expression and proliferation rate of the examined cell lines (R^2^=0.003, p=0.884).

### Elevated αSyn expression in hESC lines is compatible with cortical neuronal differentiation

Multiple clonal lines were differentiated into cortical neural progenitors using a dual-Smad inhibition protocol (Figure 2A) (Chambers et al., 2009; Shi et al., 2012). RNA was isolated and qRT-PCR used to quantify *NCAM* and *MAPT* expression at day 11 and compared to *SNCA* expression at the start of differentiation (Figure 2B). There was no correlation in day 11 *NCAM* (R^2^=0.008, p=0.761) nor day 11 *MAPT* (R^2^=0.019, p=0.461) levels relative to *SNCA* expression for the clonal lines. hESC lines with low or high αSyn expression differentiated equally well to form PAX6-positive neuroectoderm by day 12 (Figure 2C). Furthermore, by day 45, there was similar expression of early cortical markers, TBR1 and CTIP2, in low and high αSyn groups (Figure 2D). Despite the difference in αSyn levels, both groups formed neuronal networks with the same degree of efficiency based on βIII-Tubulin immunostaining (Figure 2E). This data provides evidence that αSyn expression levels do not influence neuronal induction and differentiation potential across multiple clonal hESC lines.

**Figure 2.**
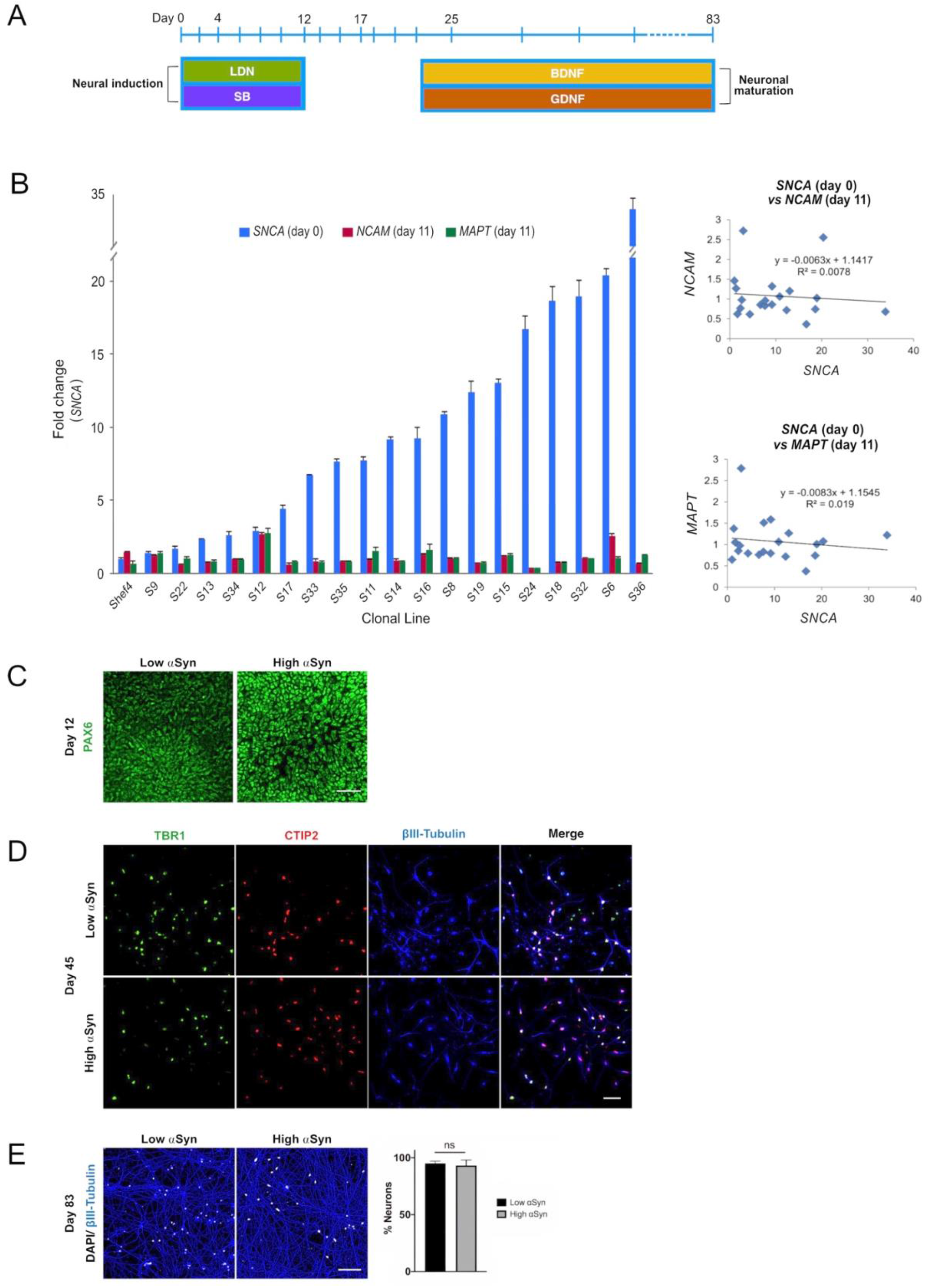
Level of αSyn does not impair cortical neuron differentiation. (A) Schematic summarising key stages of the cortical neuron differentiation protocol. Cell lifts were carried out at days 12, 17, and 25. Dual Smad inhibition using SB431542 (SB) and LDN-193189 (LDN) was employed early to drive neural induction and cortical identity, and the growth factors BDNF and GDNF were used to promote neuronal maturation after day 25. (B) Quantitative RT-PCR data measuring total *SNCA* expression in undifferentiated transgenic Shef4 cell lines, and *NCAM* and *MAPT* mRNA levels at day 11. Data shown as relative fold-change to expression in the parental Shef4 line, and error bars represent the standard error of the mean (SEM). Linear regression analysis of *SNCA, NCAM*, and *MAPT* expression levels was performed (p = 0.174 and p = 0.191 for *NCAM* and *MAPT*, respectively). (C) Representative immunocytochemistry images at day 12 of low αSyn and high αSyn hESCs differentiated into neuroectoderm, stained for the cortical progenitor marker PAX6. Scale bar = 50 μm. (D) Representative immunocytochemistry images of both low αSyn and high αSyn immature neurons (day 45) stained for the deep cortical layer markers TBR1 (green), CTIP2 (red), and βIII-Tubulin (blue), as well as merged images. Scale bar = 50 μm. (E) Immunocytochemistry images of neuronal networks representing both low αSyn and high αSyn hESCs differentiated into mature neurons (day 83), stained for the neuronal marker βIII-Tubulin (blue) and DAPI (white). Scale bar = 100 μm. Percentage neurons was calculated by quantifying DAPI levels relative to βIII-Tubulin. Significance performed using a Welch’s t-test. N = number of biological replicates (differentiated cell line) and n = technical replicates (number of wells per cell line) (N = 2, n = 5 for low αSyn; N = 3, n = 8 for high αSyn).

To determine if transgenic SNCA expression was maintained during differentiation, total and transgenic *SNCA* levels were measured by qRT-PCR (Figure 3A). The primers for total *SNCA* were designed to target the coding region and primers for transgenic *SNCA* targeted the IRES region (Figure 3A). Total and transgenic levels of human *SNCA* were measured in self-renewing hESCs (day 0) and day 25 differentiated cortical cells. Day 25 was a suitable time point as the neural induction period has been completed and immature cortical neurons are forming. Total *SNCA* expression at day 0 was significantly higher in the high αSyn vs low αSyn hESC group (Figure 3B). Importantly, at day 25, the elevated expression of *SNCA* was maintained between the high αSyn vs low αSyn cortical neuron group (p<0.01). Total *SNCA* levels did not significantly increase over each time point, day 0 vs day 25, for both high αSyn and low αSyn groups. Transgenic *SNCA* levels in undifferentiated hESCs were significantly higher in the high αSyn vs low αSyn group (Figure 3B) (p<0.01). Day 25 high αSyn vs low αSyn cortical neurons also had higher transgenic *SNCA* (p<0.01). Transgenic *SNCA* levels did not significantly increase or decrease between day 0 and day 25 for the high αSyn and low αSyn groups. Immunostaining and western blotting confirmed that high αSyn expression was maintained following cortical neuron differentiation (Figure 3C,D). At day 72 of cortical differentiation the hiPSC line, AST18, containing an *SNCA* triplication mutation exhibited a 3-fold increase in αSyn expression compared to a control hiPSC line, NAS2, derived from a 1^st^-degree relative (Devine et al., 2011) (Figure 3D). The S37 transgenic cell line maintained high αSyn expression similar to AST18 neurons, while the S36 hESC line, that expressed the most *SNCA* at day 0 (Figure 1B), now had low αSyn expression by day 72 of differentiation, suggesting significant transgene silencing (Figure 3D). This is not unexpected as transgene expression is contingent on several factors, including site of integration, number of copies of the transgene, as well as transgene silencing over cell passage or differentiation (Liu et al., 2009).

**Figure 3.**
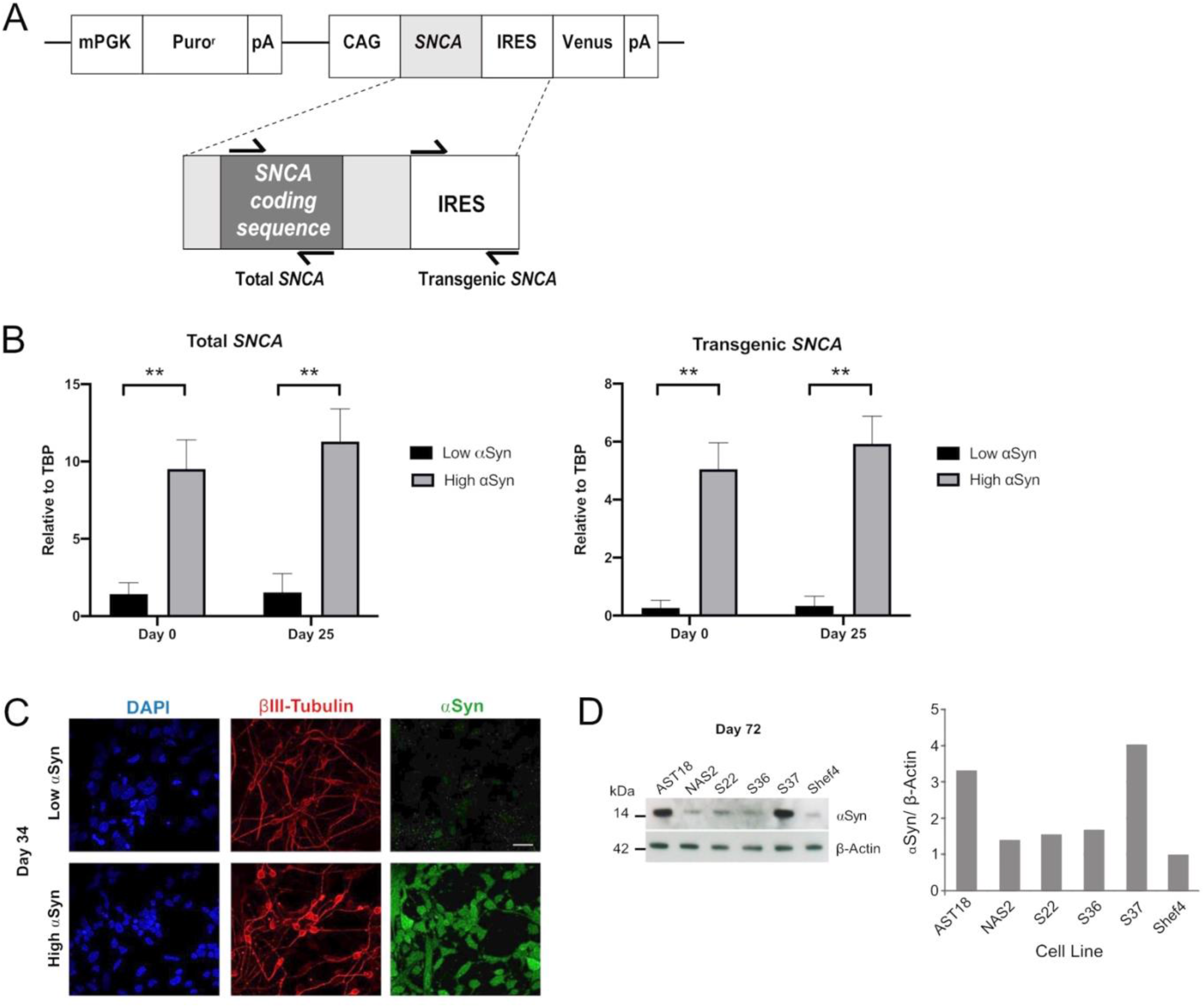
Overexpression of αSyn is maintained during cortical neuron differentiation. (A) Schematic showing the regions of *SNCA* cDNA amplified to measure “Total *SNCA*”, and location of primers targeting the internal ribosome entry site (IRES) region to measure “Transgenic *SNCA*” levels. (B) Quantitative RT-PCR for total *SNCA* and transgenic *SNCA* expression for both low αSyn and high αSyn samples at day 0 (self-renewing hESCs) and day 25 (immature differentiated cortical neurons). Mean expression levels were quantified relative to the expression of *TBP* and error bars represent SEM. N = number of biological replicates (differentiated cell line) and n = technical replicates (number of wells per cell line). For both total and transgenic αSyn, low αSyn (N = 4, n = 4) and high αSyn (N = 6 and n = 6). Statistical comparisons were performed using the Welch’s t-test (** = 2-tailed p < 0.01). (C) Representative immunocytochemistry images of differentiated neuronal cells (day 34) derived from transgenic Shef4 clonal lines S37 and S9, stained for DAPI (blue), βIII-Tubulin (red), and αSyn (green). Scale bar = 30 μm. (D) Western blot (left) for total αSyn and β-actin in cortical neurons (day 72) differentiated from transgenic hESC and hiPSC cell lines. αSyn levels quantified in ImageJ (right) for all cell lines relative to expression in Shef4-derived neurons.

### αSyn over-expression during cortical differentiation does not alter the transcriptional signature associated with neurogenesis

The transcriptomic profile of self-renewing hESC lines (day 0) and differentiated cortical neurons (day 25) with high αSyn and low αSyn were investigated using RNA-seq analysis. A total of 17 samples were subjected to UPX 3’-sequencing to measure polyadenylated mRNA transcripts across 4 sample groups (i) low αSyn hESCs, (ii) high αSyn hESCs, (iii) low αSyn cortical neurons, and (iv) high αSyn cortical neurons (Figure 4A). A Principal Component Analysis (PCA) plot showed a clear segregation between hESCs and cortical neurons, represented by 83% variance on PC1 (Figure 4B). The PC2 axis was representative of αSyn-related differences between hESC and cortical neuron samples. The hESC samples were intermixed, but cortical neuron samples segregated based on αSyn expression, however, the variance along PC2 was small (5%). This is further corroborated by hierarchical clustering analysis based on all differentially expressed genes across the four sample groups (Figure 4C).

**Figure 4.**
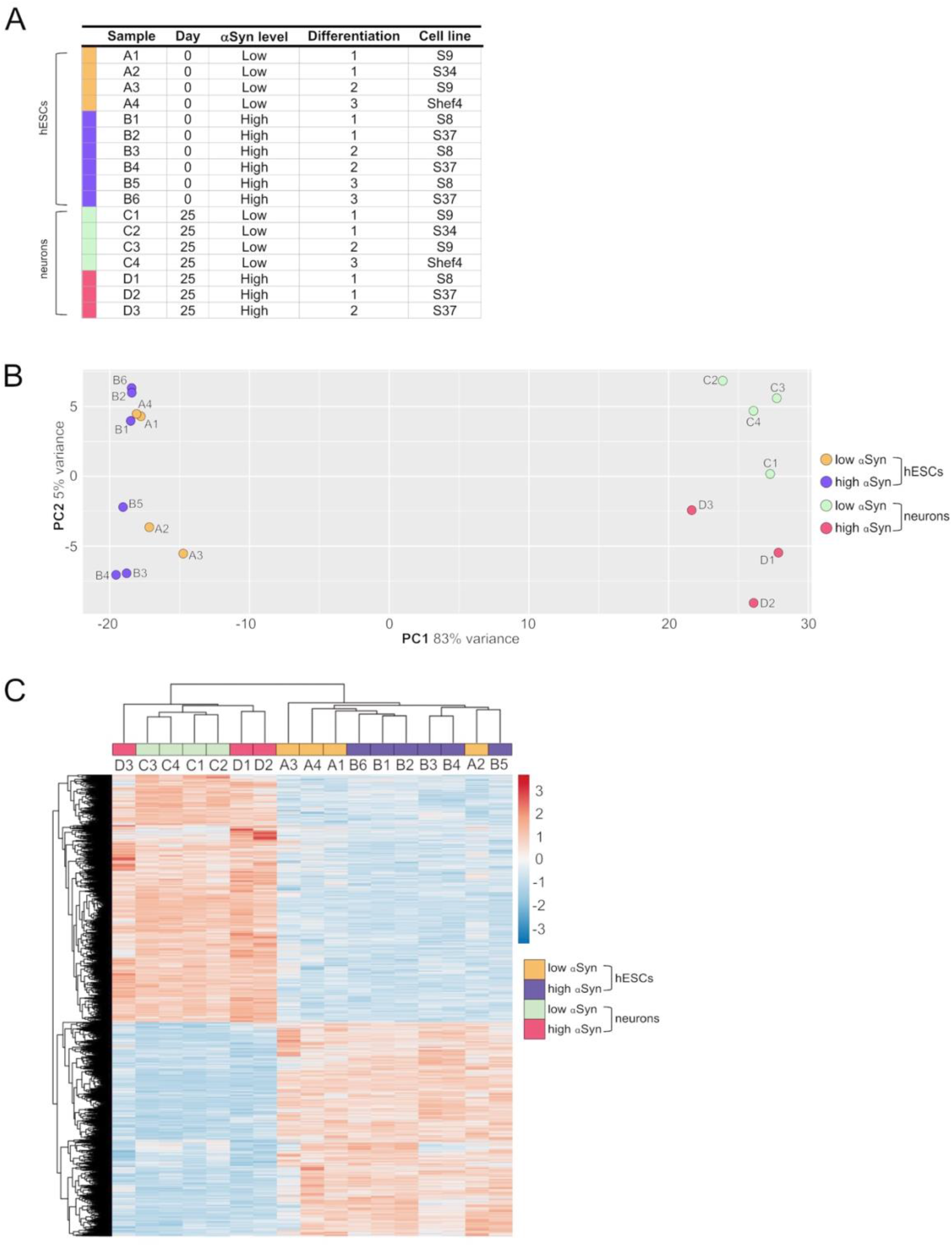
RNA-seq analysis of clonal hESC lines and differentiated cortical neurons. (A) Table of individual sample groups highlighting αSyn levels, differentiation experiment group, and the cell line of samples. N = 3 for high αSyn cortical neurons, N = 4 for both low αSyn hESCs and cortical neurons, and N = 6 for high αSyn hESCs. (B) Principal component analysis (PCA) plot of all samples. (C) Heatmap of a hierarchical cluster analysis of differential expression results, showing relative gene expression changes as either upregulated (red) or downregulated (blue). Analysis was carried out with log_2_-transformed raw counts. Plot encompasses all differentially expressed genes from all four DESeq2 pairwise comparisons, representing 5157 genes in total.

A pairwise comparison of self-renewing high αSyn vs low αSyn hESC lines found only 2 genes (*CBR1, CTNNA3*) were significantly differentially expressed between the two groups (Figure 5A), indicating αSyn does not impact on the pluripotency transcriptome. As expected, comparisons of low αSyn cortical neurons and low αSyn hESCs, as well as high αSyn cortical neurons and high αSyn hESCs, identified large numbers of differentially expressed genes (3429 and 4188, respectively, Figure 5B,C; Supplementary Table S1). Seven of the top ten most significantly upregulated genes were the same for these two pairwise comparisons, including *SOX5, NPAS3, MALAT1, MAP2, QKI, PCDH9* and *AC10729.1* (Figure 5D). Most of these genes have a role in neurogenesis (Brunskill et al., 1999; Chen et al., 2016; Hardy et al., 1996; Izant and McIntosh, 1980; Strehl et al., 1998; Wunderle et al., 1996). Similarly, five of the top ten significantly downregulated genes were the same in the hESCs *vs* cortical neuron comparisons for high and low αSyn cell lines, including *DPPA4, L1TD1, RBM47, XACT* and *DNMT3B* (Figure 5D). Most of these top downregulated genes have roles in pluripotency (Emani et al., 2015; Hu et al., 2012; Madan et al., 2009; Radine et al., 2020; Vallot et al., 2013). There was a large overlap in the genes that were significantly upregulated and downregulated in cortical neuron vs hESC groups with high or low αSyn (Figure 6A). There was also a substantial overlap of gene ontology (GO) terms significantly upregulated and downregulated in the high or low αSyn comparisons (Figure 6B). The most significantly enriched KEGG pathway in both high and low αSyn cortical neuron vs hESC comparisons for upregulated genes was ‘axon guidance’ (KEGG:04360). Five other top ten KEGG pathways linked with upregulated genes were the same for both the high αSyn and low αSyn cortical neuron vs hESC comparisons (Figure 6C). The ‘ribosome’ term (KEGG:03010) was the most significant downregulated pathway common to both comparisons. Interestingly, the Parkinson’s disease KEGG pathway was downregulated in high αSyn cortical neurons vs high αSyn hESCs, due to the downregulation of several mitochondrial genes and cytochrome c oxidase genes, including *NDUFB9, MT-CO2, MT-CO3, MT-ATP6, MT-ATP8, MT-CYB, COX7C* and *CYCS*. This may reflect a mitochondrial phenotype caused by elevated αSyn expression in cortical neurons. The data so far suggests that the process of cortical neuron differentiation induced a large number of significant gene expression changes, and that the level of αSyn in these cells does not impair this process.

**Figure 5.**
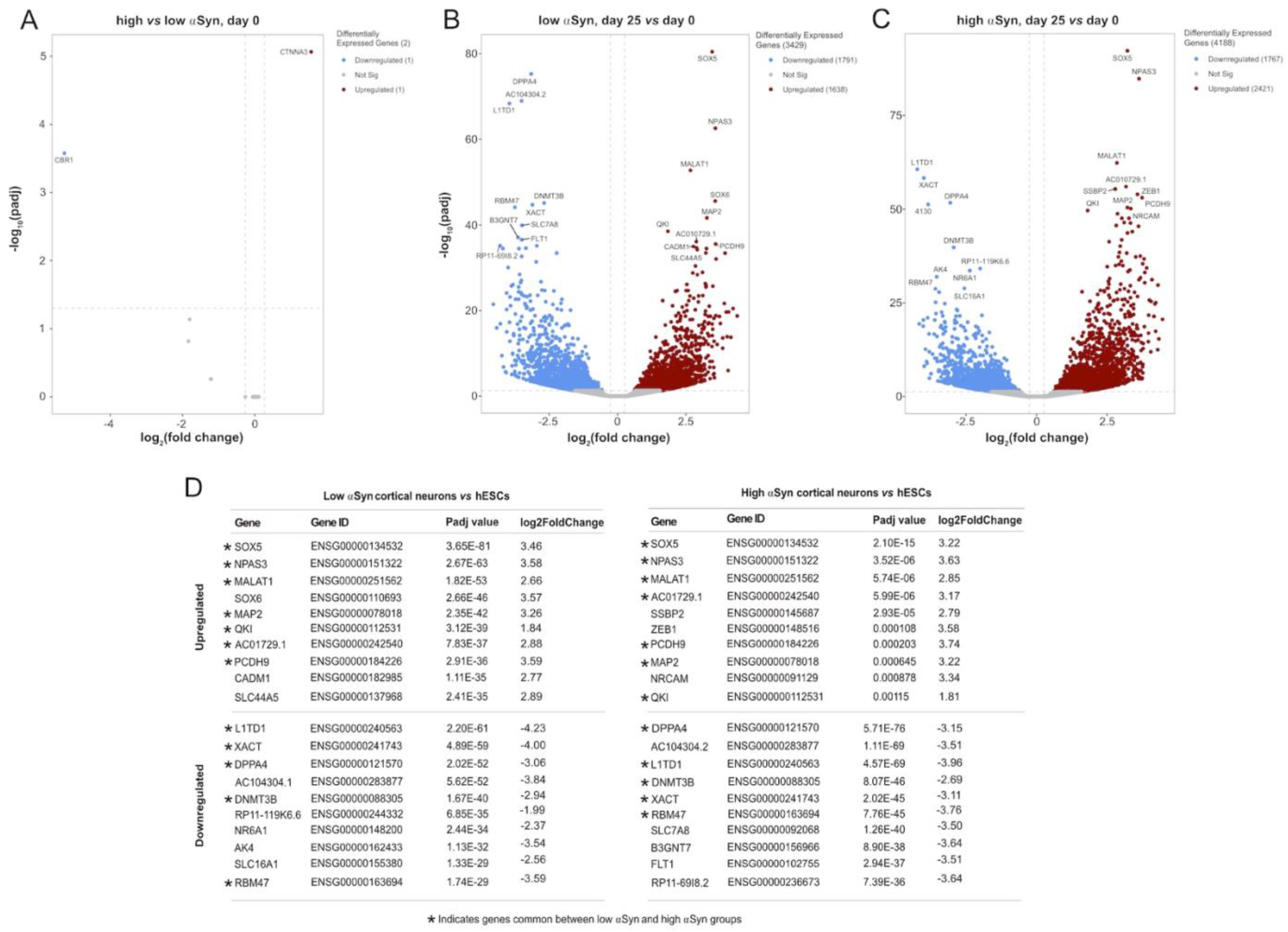
RNA-seq reveals large and over-lapping gene expression changes between cortical neurons and hESCs for both low αSyn and high αSyn groups. (A-C) Volcano plots showing differentially expressed genes between (A) low αSyn hESCs and high αSyn hESCs, (B) low αSyn cortical neurons (day 25) *vs* hESCs (day 0), and (C) high αSyn cortical neurons (day 25) *vs* hESCs (day 0). Horizontal dashed lines cross the y-axis at –log_10_(0.05), representing a significance cut-off Padj value of 0.05. Vertical lines represent a fold-change cut-off of 1.2, and therefore cross the x-axis at 0.263 and - 0.263.

**Figure 6.**
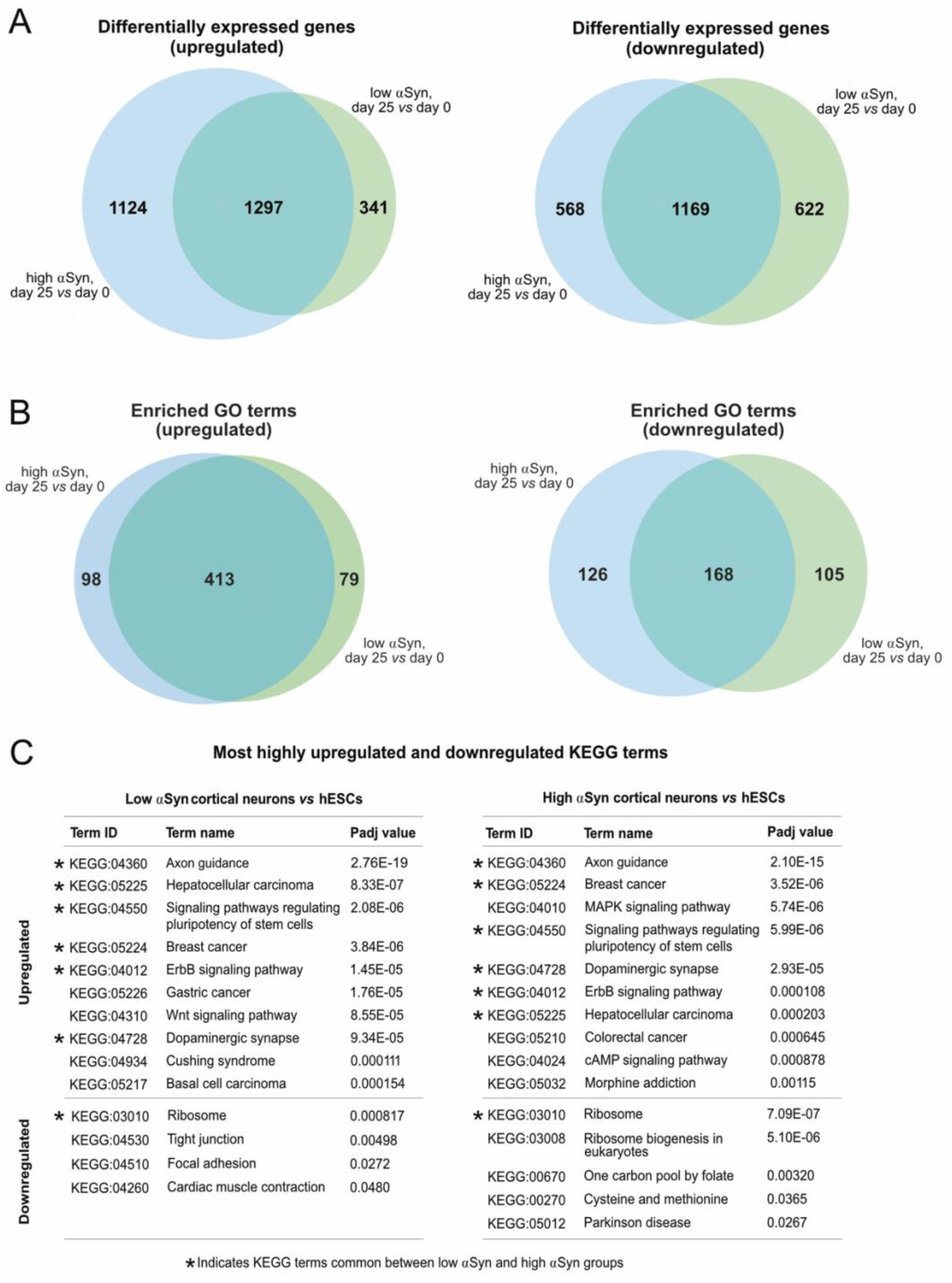
(A) Venn diagrams comparing upregulated and downregulated differentially expressed genes from low αSyn cortical neurons *vs* hESCs and high αSyn cortical neurons *vs* hESCs comparisons. (B) Venn diagrams comparing upregulated and downregulated GO terms from low αSyn cortical neurons *vs* hESCs and high αSyn cortical neurons *vs* hESCs comparisons. (C) List of top 10 upregulated and top 10 downregulated KEGG terms by Padj value, in order of ascending Padj, for low αSyn cortical neurons *vs* hESCs and high αSyn cortical neurons *vs* hESCs (if less than 10, all terms shown). KEGG terms common to both comparisons are indicated.

The key group comparison was that of high αSyn cortical neurons vs low αSyn cortical neurons. The volcano plot, in comparison to the cortical neurons vs hESC plots, identified a total of 47 differentially expressed genes (Figure 7A; Supplementary Table S1). KEGG pathway analysis did not show any relevant neurogenesis-related pathways enriched in high αSyn vs low αSyn cortical neurons (Figure 7B). Selected gene expression analysis revealed that the pluripotency markers *POU5F1 (OCT4)* and *NANOG* were significantly downregulated and neurogenesis markers, *ASCL1, MYT1L*, and *POU3F2*, were significantly upregulated in cortical neuron samples relative to hESC samples for both high αSyn and low αSyn groups (Figure 7C) (Chambers et al., 2003; Nichols et al., 1998; Vierbuchen et al., 2010). Analysis of genes implicated in axon guidance, including *MAP2, SLIT2, EPHB1* and *NCAM1* showed no difference in cortical neurons with low or high αSyn, except for *NTN1* (p<0.01) (Figure 7D) (Borrell et al., 2012; Enriquez-Barreto et al., 2012; Izant and McIntosh, 1980; Kennedy et al., 1994). There were no significant differences in genes associated with synaptic development, in low αSyn and high αSyn cortical neurons, including *CADM1, NLGN1, SNAP25, SYP* and *DLG4* (Figure 7E) (Cho et al., 1992; Ichtchenko et al., 1995; Oyler et al., 1989; Stagi et al., 2010; Wiedenmann and Franke, 1985). Other markers were explored to ascertain telencephalic development. *FOXG1* and *PAX6*, cortical progenitor markers, and *CTIP2* and *TLE4*, markers of deep cortical layers, showed no significant differences in low and high αSyn cortical neuron groups (Figure 7F) (Raciti et al., 2013). The superficial layer marker *SATB2*, was not significantly different between the low and high αSyn cortical neurons, which is in keeping with its involvement in the later stages of cortical development (Mariani et al., 2012). Expression of the neuronal migration marker, *DCX*, also presented no significant variation between low or high αSyn cortical neurons (Figure 7F) (Gleeson et al., 1999).

**Figure 7.**
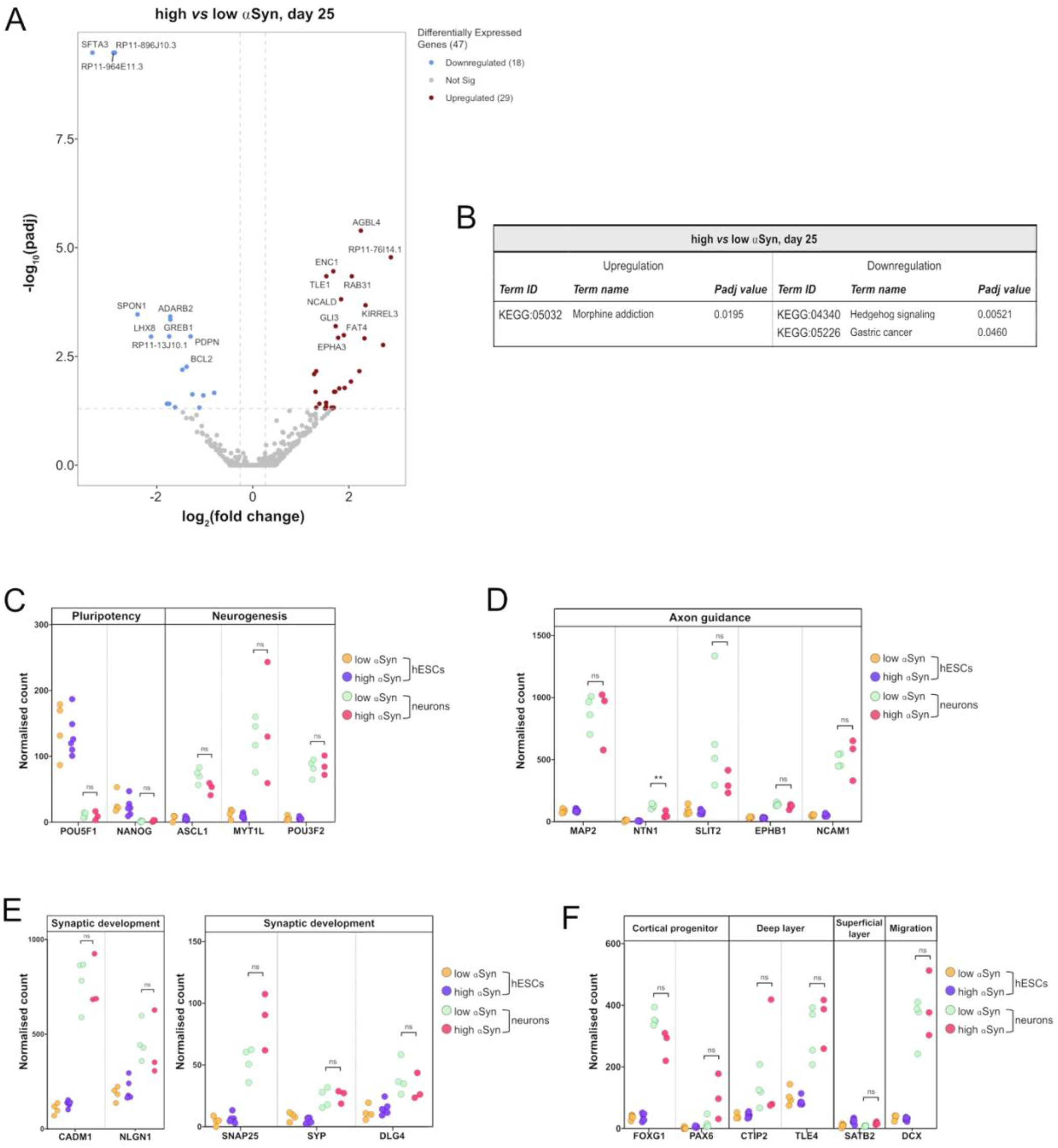
RNA-seq analysis reveals that αSyn overexpression does not have a significant impact on cortical marker gene expression. (A) Volcano plot showing distribution of differentially expressed genes between high αSyn and low αSyn day 25 cortical neurons (B) Table showing all upregulated and downregulated KEGG terms for this comparison. (C-F) Plots showing normalised single gene read counts across all four sample groups for (C) pluripotency and neurogenesis markers, (D) axon guidance genes, (E) synaptic development genes, and (F) cortical progenitor, deep layer, superficial layer, and migration genes. Statistical comparisons between cortical neurons were performed using the Welch’s t-test (** p < 0.01).

## Discussion

In this study, we generated and investigated 29 *SNCA* transgenic hESC lines and found no correlation between αSyn expression level and neural differentiation potential. We further addressed the hypothesis that increased αSyn expression does not impair cortical neuronal differentiation of human pluripotent stem cells. Across multiple clonal lines, multiple rounds of cortical neuron differentiation using two robust differentiation protocols and unbiased transcriptomic analysis, we show data to support this hypothesis. The level of *SNCA* expression, whether normal or increased, did not have a significant impact on the efficiency of cortical neuron differentiation. These findings resolve conflicting data in the field, and are highly relevant to studies utilising human pluripotent stem cells (hESCs or iPSCs) with differential αSyn expression to model synucleinopathies, such as PD, DLB and multiple system atrophy (MSA).

As expected, significant differential gene expression was observed when cortical neurons were compared to undifferentiated hESCs for both high αSyn and low αSyn hESCs. Gene ontology (GO) and KEGG pathway analysis identified significant overlap in the group comparisons with high αSyn or low αSyn hESCs and their differentiated counterparts. Furthermore, most of the top differentially expressed genes were the same in these comparisons, indicating neurogenesis proceeded in a similar manner despite the level of αSyn expression. The gene expression differences that were unique to the high αSyn hESCs or to the low αSyn hESCs could be due to a number of factors, including clonal variation, experiment-to-experiment variation, or due to a phenotype of elevated αSyn expression, such as mitochondrial dysfunction. When high αSyn and low αSyn cortical neurons were directly compared to each other, only 47 genes were differentially expressed, and pathway analysis did not reveal anything relevant to cortical neuron induction or differentiation. Furthermore, read count analysis for well-characterised genes with known roles in neurogenesis and the development of cortical identity confirmed cortical differentiation occurred similarly between the high αSyn and low αSyn groups.

Research groups, including ourselves, have used hiPSCs derived from patients with *SNCA* multiplication mutations to model synucleinopathies (Brazdis et al., 2020; Byers et al., 2011; Chen et al., 2019; Devine et al., 2011; Flierl et al., 2014; Lin et al., 2016; Oliveira et al., 2015; Prots et al., 2018). We previously showed, using eight clonal hiPSC lines, harbouring the *SNCA* triplication mutation and six hiPSC lines from a non-affected first-degree relative, that the main sources of variation in differentiation efficiency were due to differences between clonal lines, reprogramming efficiency, and the process of neuronal differentiation itself (Devine et al., 2011). Whilst cellular reprogramming is a powerful method for generating human disease models, reprogramming can be incomplete and can introduce coding mutations or large chromosomal abnormalities that may lead to altered differentiation potential of different clones (Boulting et al., 2011; Hussein et al., 2011; Mayshar et al., 2010). Furthermore, in these patient-derived iPSCs the size of the multiplicated region is variable and other adjacent genes such as *MMRN1* may be incorporated (Ross et al., 2008). Over-expression of other coding genes in these iPSC lines presents another potential confounding factor when investigating and interpreting neuronal differentiation potential. A number of these caveats apply to studies differentiating hiPSCs with a *SNCA* triplication into neuronal progenitors or dopaminergic neurons (Flierl et al., 2014; Oliveira et al., 2015). In particular, limited clonal lines were examined and siRNA knock-down of *SNCA* ‘rescue’ of dopaminergic differentiation was partial (Oliveira et al., 2015).

To further highlight the point regarding clonal variation and conflicting reports in the literature, recent studies have shown, using hiPSCs from patients with a *SNCA* duplication mutation, that cortical neurons (Prots et al., 2018) and dopaminergic neurons can be efficiently and comparably generated using these hiPSC lines (Brazdis et al., 2020). Other independent groups have also used hiPSCs with a *SNCA* triplication mutation to show functional dopaminergic neuron generation comparable to control hiPSC lines (Byers et al., 2011; Lin et al., 2016). We recently showed that reducing *SNCA* alleles in isogenic hESC lines also does not affect neuronal differentiation; wild type, *SNCA*^*+/-*^ and *SNCA*^*-/-*^ hESC lines showed no differences in differentiation into dopaminergic neurons (Chen et al., 2019). In the studies reporting no impairment of dopaminergic neuron differentiation with a *SNCA* triplication, PD-related phenotypes, including reduced synchronous firing on microelectrode recordings and increased susceptibility to oxidative stress were observed (Brazdis et al., 2020; Lin et al., 2016). While there may be αSyn-related phenotypic differences or inherent vulnerabilities due to elevated αSyn expression in mature neurons, this does not imply that the process of neuronal differentiation itself is impaired.

When αSyn was over-expressed using a lentiviral system in hESC-derived neuroectoderm impaired neuronal patterning and acute toxicity were reported (Schneider et al., 2007). Dopaminergic and GABAergic neuron populations were affected by αSyn over-expression, but this was only quantified in a single hESC line (H9), and the toxicity shown to be occurring could result in selective neuronal loss, making the interpretation of differentiation marker analysis challenging (Schneider et al., 2007). Similar caveats apply to work performed using AF22 neural stem cells with doxycycline-inducible αSyn expression. Exposure to doxycycline may cause pleotropic effects, and αSyn expression was not sustained throughout differentiation (Zasso et al., 2018).

It is important to consider that neuronal differentiation protocols often require optimisation for each clonal line and small variations in ligand concentrations, specifically CHIR99021 in the midbrain dopaminergic differentiation protocol, can impair neurogenesis. For example, in the study by Oliveira *et al*., the dopaminergic differentiation protocol yielded less than eight per cent of TH-positive neurons in the control lines (Oliveira et al., 2015). Whilst our study does not yield insight into the impact of elevated αSyn expression on dopaminergic differentiation, a subject for future studies, the cortical neuron protocols used were robust with the vast majority of cells producing cortical neurons.

In summary, this work has produced an allelic series of isogenic clonal hESC lines with differing levels of αSyn expression, and we have demonstrated that elevated αSyn expression does not impair cortical neurogenesis. This supports the validity of using human pluripotent stem cells, such as iPSCs with *SNCA* multiplications and transgenic hESC lines, to model synucleinopathies, and highlights the need for efforts to closely match the differentiation stage and maturity of neurons prior to commencing with disease modelling.

## Methods

### Generation of transgenic hESC lines by nucleofection

Shef4 hESCs were provided Prof D Hay (University of Edinburgh) following MRC Steering Committee approval (SCSC11-60). The plasmid consisting of wild-type human *SNCA* (pcDNA3.1) was provided by Prof J Hardy (UCL). The PGK-Puro-pCAGS and FCT-IRES-Venus-pBS plasmids were provided by William Hamilton (University of Edinburgh).

IRES-Venus and human *SNCA* fragments were amplified by PCR (MJ Research, PTC-200 Peltier Thermal Cycler) and purified using DNA Clean & Concentrator™-5 Kit (Zymo Research, D4003) to provide 20 μl of eluted DNA. 1X BSA, 1X digestion buffer and 50 U digestion enzyme (all New England Biolabs®) were used to digest the plasmid DNA, which was then purified from agarose gel using Zymoclean™ Gel DNA Recovery Kit (Zymo Research, D4001) per manufacturer’s protocol. Purified and digested DNA was ligated to PCR products at 16°C overnight in a final volume of 20 μl using 2 U of T4 DNA ligase and 1x ligation buffer (Roche, 10481220001). Plasmid DNA was transformed into TOP10 chemically competent cells (Invitrogen, c4040-10) by the heat-shock method. Plasmid DNA was extracted using QIAprep^®^-Spin Miniprep kit (Qiagen, 27104) or Maxiprep (Qiagen, 12662) systems per manufacturer’s protocol, then desalted using Millipore centrifugal filter units (Millipore, UFC503024 24PK). The final pCAG-SNCA-IRES-Venus construct contained human *SNCA*, internal ribosome entry site (IRES) and Venus expression cassette under the constitutive pCAG promoter, as well as a puromycin resistance gene (Puro^r^) driven by the PGK promoter. The control construct, pCAG-IRES-Venus, contained the same elements except for the *SNCA* gene (Figure 1A).

The Neon Transfection System (Invitrogen, MPK5000) was used for nucleofection of both constructs into Shef4 hESCs as per the manufacturer’s protocol. 1 μg/ml puromycin was added to the culture media for at least two weeks to isolate clones and colonies, manually picked for expansion and cryopreservation. Clones were screened for *SNCA* over-expression by qRT-PCR.

### Quantitative RT-PCR

The MasterPure™ Complete DNA and RNA Purification kit (Epicentre, MC85200) or the RNeasy kit (Qiagen, 74104) was used for RNA extraction. Genomic DNA was removed using DNase I (Promega, M6101). cDNA was synthesised from 500 ng total RNA using M-MLV reverse transcriptase (RT, ThermoFisher Scientific, 28025013) or Superscript IV reverse transcriptase (Invitrogen, 18090010). qRT-PCR was performed using a LightCycler™ 480 (Roche) with the following parameters: (95°C for 10 min), [(95°C for 10 sec) + (60°C for 20 sec)] over 45 cycles. Intron-spanning primers were designed using the Universal Probe Library (UPL) Assay design centre (Roche). Primer sequences and UPL probes were total *SNCA* F-tgggcaagaatgaagaaggagc, R-gtggtgacgggtgtgacagc Probe 68; transgenic *SNCA* F-cgacctgcagttggacct, R-tgacaatgacatccactttgc Probe 163; *NCAM* F-gcgttggagagtccaaattc, R-gggagaaccaggagatgtcttt Probe 51; *MAPT* F-accacagccaccttctcct, R-cagccatcctggttcaaagt Probe 55; TATA-box binding protein (*TBP*) F-atagggattccgggagtcat, R-gaacatcatggatcagaacaaca Probe 87. Each 10μl reaction was performed in triplicate and results normalised to *TBP* expression.

### MTS Assay

Cell proliferation was assessed using a colorimetric assay, CellTiter96^®^ AQ_ueous_ One Solution Cell Proliferation Assay or MTS assay. In this test, MTS tetrazolium is bioreduced in viable cells (Supplementary Figure 1).

### hESC and hiPSC culture

Shef4-derived transgenic hESC lines, and AST18 and NAS2 hiPSCs were expanded in culture on either Matrigel-coated 6-well plates (BD, 356234) in mTeSR1 medium (Stemcell™ Technologies, 05850) or on Laminin-521 (BioLamina, LN521) coated 6-well plates in StemMACS iPS-Brew XF, human (Miltenyi Biotec, 130-107-086). 1 μg/ml puromycin (Sigma, P8833) was used in the culture media of Tg hESCs to maintain the expression of the transgenes.

### Cortical neuronal differentiation

Two cortical neuron differentiation protocols were used, referred to as CD protocol 1 and CD protocol 2, respectively. CD protocol 1 was adapted from the cortical neuron differentiation protocol published by Chambers et al., in 2009 and CD protocol 2 was adapted from the published protocol by Shi et al., in 2012 (Chambers et al., 2009; Shi et al., 2012). Both protocols use dual Smad inhibition to induce cortical neuron differentiation. In CD protocol 1, neural differentiation was started by changing the culture media to neural induction media (NIM) which included 10 μM SB431542 (Tocris, 616461) and 100 nM LDN-193189 (Stemgent, 04-0019). NIM for this protocol was prepared by mixing 1:1 DMEM/F12 (Gibco, 20331-020) and Neurobasal medium (Gibco, 21103-049), supplemented with 1 ml N2 and 2 ml B27 with retinoic acid (Gibco, 17504-044). NIM was also supplemented with 2 mM L-glutamine, 0.1 mM β-mercaptoethanol (BDH, 44143-31), 100 U/ml penicillin and 100 μg/ml streptomycin (Invitrogen, 15140-122) and 100μM non-essential amino acids (Gibco, 1140-035). Cells were lifted using dispase at day 12, dissociated into clumps and plated on 10 μg/ml Laminin-111 (Sigma L2020-1MG) and 15 μg/ml poly-L-ornithine-coated plates (Sigma, P4957). 100 nM LDN and 20 μg/ml FGF2 were included in the NIM, and SB431542 removed from day 12. Following 7-10 days of progenitor colony expansion, cells were lifted using accutase and plated onto Laminin-111/ poly-L-ornithine-coated plates in NIM supplemented with 10 ng/ml BDNF (Peprotech, 450-02) and 10 ng/ml GDNF (Peprotech, 450-10). Half media changes were performed every 3 days during the neuronal maturation phase up to day 83.

In CD protocol 2 hESCs at 80%-90% confluency in 6-well plates were lifted with 1 ml/well UltraPure 0.5 M EDTA (Invitrogen, 15575038), counted, and transferred onto 5 μg/ml Laminin-111 coated (Biolamina, LN111-04) 24-well plates (Corning, 3527), at an initial plating density of 80,000 cells/cm^2^. 600 μl/well neural induction media (NIM) was used until day 4 of differentiation. NIM was composed of 50% DMEM/F12 (ThermoFisher Scientific, 21331020) and 50% Neurobasal Media (ThermoFisher Scientific, 21103049), B27 supplement with Retinoic Acid (ThermoFisher Scientific, 17504044), N2 supplement (ThermoFisher Scientific, 17502048) and 2 mM L-Glutamine (ThermoFisher Scientific, 25030123). From day 4 onwards, 50% NIM, 25% DMEM/F12 and 25% Neurobasal Media with 2 mM L-glutamine. For the first 12 days of differentiation, 10 μM SB431542 (Tocris, 616461) and 100 nM LDN-193189 (Miltenyi Biotec, 130-103-925) were added to the NIM and this media was replaced every two days. Cells were lifted at day 12 and day 17 with Collagenase Type IV (Life Technologies, 17104019) diluted in HBSS (ThermoFisher Scientific, 14025). Cell were lifted and passaged as clumps in a ratio of 1:1.5 and 1:2 at day 12 and 17, respectively. 10 μM Y27632 (Tocris) was used in the media for each cell lift. At day 25, differentiated cells were lifted with Accutase (Sigma, A6964) and cell pellets frozen using a dry ice and ethanol bath for RNA isolation.

### Transcriptomic analysis

Total RNA was isolated with RNeasy kit (QIAGEN, 74104). RNA integrity (RINe ≥ 7) was confirmed using Tapestation 4200 (Agilent). The median RINe score was 9.2. Samples were collected at day 0 and day 25, across three sets of cortical neuron differentiations (Figure 4A). The samples were processed by Qiagen Genomic Services using their QIAseq UPX 3’ Transcriptome kit and libraries were sequenced on a NextSeq500 instrument. The total number of polyadenylated 3’ transcript reads and the mean number of reads per unique molecular identifier were counted. The raw transcript counts were analysed using DESeq2 (Love et al., 2014) differential expression analysis in R studio. Pairwise analysis was used to compare hESC vs cortical neurons with high or low αSyn. A padj cut off value of 0.05 and a log_2_ fold-change cut off value of 1.2, were used. KEGG pathway analysis was performed using the g:GOSt function in the CRAN gprofiler2 package (Raudvere et al., 2019). RNA-seq data has been deposited on the Gene Expression Omnibus (GEO Accession number: GSE195877).

### Immunocytochemistry

Immunostaining was performed on cells cultured on 13-mm glass coverslips or in Ibidi 8-well plates. Spent medium was removed and cells fixed with 4% PFA for 15 minutes. Following 3 PBS (ThermoFisher Scientific) washes, the cells were permeabilised and blocked with 2% goat (or donkey) serum (Sigma) in 0.1% Triton-X-100 (Fisher) in PBS for 45 minutes prior to overnight incubation at 4°C with primary antibodies. The primary antibodies used were β-III tubulin (1:1,000, mouse IgG2b, Sigma T8660), CTIP2 (1:500, rat IgG2a, Abcam ab18465), PAX6 (1:40, mouse IgG1, DSHB ab528427), TBR1 (1:200, rabbit IgG, Abcam ab31940) and total αSyn (1:1000, mouse IgG1, BD 610787). Secondary antibodies were applied at room temperature for 1 hour in the dark. These were Alexa Fluor 488 (ThermoFisher Scientific, A21121), Alexa Fluor 488 (ThermoFisher Scientific, A21131), Alexa Fluor 488 (ThermoFisher Scientific, A21208), Alexa Fluor 555 (ThermoFisher Scientific, A21127), Alexa Fluor 555 (ThermoFisher Scientific, A21428), Alexa Fluor 555 (ThermoFisher Scientific, A31572), Alexa Fluor 647 (ThermoFisher Scientific, A21242) and Alexa Fluor 647 (ThermoFisher Scientific, Ab150107). Following a further three PBS washes, slides were mounted using Fluorsave (Merck, 345789). 0.1 μg/ml 4′,6-diamidino-2-phenylindole (DAPI, Life Technologies) was used to stain nuclei.

### Image capture, processing and quantification

The Eclipse Ti (Nikon) and/ or the Axio Observer (Zeiss) microscopes were used to acquire the images presented. Huygens Software (Scientific Volume Imaging) was used for deconvolution of Z stack images (at least 10 images/ stack and maximum intensity pixel projection used). Fiji software was used for image analysis and quantification, with identical brightness and contrast values used for each image channel and for each experiment. Macro scripts were used to split the image channels, threshold and binarize for quantification.

### Western blotting

RIPA Lysis Buffer (Santa Cruz, sc-24948) was used to lyse cell pellets and protein concentration determined using the Micro BCA Protein Assay kit (ThermoFisher Scientific, 232350). 10 μg protein, per sample, was mixed and incubated with 5 μl NuPAGE™ LDS Sample Buffer (ThermoFisher Scientific, NP0007) and 2 μl 1M DTT (ThermoFisher Scientific, NP0004) prior to loading on a NuPAGE™ 4%–12% Bis-Tris Gradient Gel (ThermoFisher Scientific, NP0322BOX). SeeBlue™ Plus2 Pre-stained Protein Standard (5 μl, Thermo Fisher Scientific, LC5925) was used as a standard. Following electrophoresis, protein was transferred onto a 0.45 μm nitrocellulose (Amersham Protran Premium, 10600096) or a PVDF membrane (GE Healthcare Amersham Hybond ECL, RPN68D). 0.4% PFA was used to fix the protein on the membrane. One minute of methanol (Fisher Scientific, M/3900/17) immersion was done, in addition, if a PVDF membrane was used. Membranes were immersed in blotting-grade blocker (BioRad, 1706404) in 0.1% TBS-Tween for an hour at room temperature and then the primary antibody, mouse anti-αSyn (1:1000, BD) overnight at 4^°^C. Following three washes in 0.1% TBS-Tween, 1:2000 HRP-conjugated anti-mouse IgG (Promega) secondary antibody was applied for two hours at room temperature. Pierce™ ECL Western Blotting Substrate (ThermoFisher Scientific, 32109) was then added to the membrane for image capture using the LI-COR Odyssey imaging system (Biosciences). Antibodies were stripped with the Restore™ PLUS Western Blot Stripping Buffer (ThermoFisher Scientific, 46430). The blocking step was repeated and secondary antibody, HRP-conjugated anti-ß-Actin antibody (1:1000, Abcam) applied. The Pierce™ ECL Western Blotting Substrate was again used prior to imaging the membrane.

### Statistics

Statistical tests were performed using SPSS v23 and include the Student’s t-test or t-test with Welch’s correction, linear regression analysis and the Mann-Whitney U test for non-parametric data. Each Figure legend details which test was used for each statistical comparison. Significance level cut off was p<0.05.

## Acknowledgments

The authors with to thank the Wellcome Trust and the MRC for funding this project. We are very grateful to Prof Steven Pollard for critical comments on the manuscript, and Dr James Ashmore for bioinformatics guidance. We also sincerely thank Prof John Hardy and Dr William Hamilton for pcDNA3.1-SNCA and PGK-Puro-pCAGS plasmids, respectively, and Prof David Hay for Shef4 hESCs.

## Author contributions

TK conceived the project. TK, AN, RY designed the experiments. AN and RY performed the experiments and analysed the data. RB performed bioinformatic analysis. AN and TK wrote the manuscript, and all authors reviewed and approved the manuscript.

## References

Abeliovich, A., Schmitz, Y., Fariñas, I., Choi-Lundberg, D., Ho, W.H., Castillo, P.E., Shinsky, N., Verdugo, J.M., Armanini, M., Ryan, A., Hynes, M., Phillips, H., Sulzer, D., Rosenthal, A., 2000. Mice lacking alpha-synuclein display functional deficits in the nigrostriatal dopamine system. Neuron 25, 239– 252. doi:10.1016/s0896-6273(00)80886-7

Al-Wandi, A., Ninkina, N., Millership, S., Williamson, S.J.M., Jones, P.A., Buchman, V.L., 2010. Absence of alpha-synuclein affects dopamine metabolism and synaptic markers in the striatum of aging mice. Neurobiol. Aging 31, 796–804. doi:10.1016/j.neurobiolaging.2008.11.001

Anwar, S., Peters, O., Millership, S., Ninkina, N., Doig, N., Connor-Robson, N., Threlfell, S., Kooner, G., Deacon, R.M., Bannerman, D.M., Bolam, J.P., Chandra, S.S., Cragg, S.J., Wade-Martins, R., Buchman, V.L., 2011. Functional alterations to the nigrostriatal system in mice lacking all three members of the synuclein family. J. Neurosci. 31, 7264–7274. doi:10.1523/JNEUROSCI.6194-10.2011

Borrell, V., Cárdenas, A., Ciceri, G., Galcerán, J., Flames, N., Pla, R., Nóbrega-Pereira, S., García-Frigola, C., Peregrín, S., Zhao, Z., Ma, L., Tessier-Lavigne, M., Marín, O., 2012. Slit/Robo signaling modulates the proliferation of central nervous system progenitors. Neuron 76, 338–352. doi:10.1016/j.neuron.2012.08.003

Boulting, G.L., Kiskinis, E., Croft, G.F., Amoroso, M.W., Oakley, D.H., Wainger, B.J., Williams, D.J., Kahler, D.J., Yamaki, M., Davidow, L., Rodolfa, C.T., Dimos, J.T., Mikkilineni, S., MacDermott, A.B., Woolf, C.J., Henderson, C.E., Wichterle, H., Eggan, K., 2011. A functionally characterized test set of human induced pluripotent stem cells. Nat. Biotechnol. 29, 279–286. doi:10.1038/nbt.1783

Brazdis, R.-M., Alecu, J.E., Marsch, D., Dahms, A., Simmnacher, K., Lörentz, S., Brendler, A., Schneider, Y., Marxreiter, F., Roybon, L., Winner, B., Xiang, W., Prots, I., 2020. Demonstration of brain region-specific neuronal vulnerability in human iPSC-based model of familial Parkinson’s disease. Hum. Mol. Genet. 29, 1180–1191. doi:10.1093/hmg/ddaa039

Brunskill, E.W., Witte, D.P., Shreiner, A.B., Potter, S.S., 1999. Characterization of npas3, a novel basic helix-loop-helix PAS gene expressed in the developing mouse nervous system. Mech. Dev. 88, 237– 241. doi:10.1016/S0925-4773(99)00182-3

Byers, B., Cord, B., Nguyen, H.N., Schüle, B., Fenno, L., Lee, P.C., Deisseroth, K., Langston, J.W., Pera, R.R., Palmer, T.D., 2011. SNCA triplication Parkinson’s patient’s iPSC-derived DA neurons accumulate α-synuclein and are susceptible to oxidative stress. PLoS One 6, e26159. doi:10.1371/journal.pone.0026159

Chambers, I., Colby, D., Robertson, M., Nichols, J., Lee, S., Tweedie, S., Smith, A., 2003. Functional expression cloning of Nanog, a pluripotency sustaining factor in embryonic stem cells. Cell 113, 643–655. doi:10.1016/s0092-8674(03)00392-1

Chambers, S.M., Fasano, C.A., Papapetrou, E.P., Tomishima, M., Sadelain, M., Studer, L., 2009. Highly efficient neural conversion of human ES and iPS cells by dual inhibition of SMAD signaling. Nat. Biotechnol. 27, 275–280. doi:10.1038/nbt.1529

Chartier-Harlin, M.-C., Kachergus, J., Roumier, C., Mouroux, V., Douay, X., Lincoln, S., Levecque, C., Larvor, L., Andrieux, J., Hulihan, M., Waucquier, N., Defebvre, L., Amouyel, P., Farrer, M., Destée, A., 2004. Alpha-synuclein locus duplication as a cause of familial Parkinson’s disease. Lancet 364, 1167–1169. doi:10.1016/S0140-6736(04)17103-1

Chen, L., Feng, P., Zhu, X., He, S., Duan, J., Zhou, D., 2016. Long non-coding RNA Malat1 promotes neurite outgrowth through activation of ERK/MAPK signalling pathway in N2a cells. J. Cell Mol. Med. 20, 2102–2110. doi:10.1111/jcmm.12904

Chen, Y., Dolt, K.S., Kriek, M., Baker, T., Downey, P., Drummond, N.J., Canham, M.A., Natalwala, A., Rosser, S., Kunath, T., 2019. Engineering synucleinopathy-resistant human dopaminergic neurons by CRISPR-mediated deletion of the SNCA gene. Eur. J. Neurosci. 49, 510–524. doi:10.1111/ejn.14286

Cho, K.O., Hunt, C.A., Kennedy, M.B., 1992. The rat brain postsynaptic density fraction contains a homolog of the Drosophila discs-large tumor suppressor protein. Neuron 9, 929–942. doi:10.1016/0896-6273(92)90245-9

Connor-Robson, N., Peters, O.M., Millership, S., Ninkina, N., Buchman, V.L., 2016. Combinational losses of synucleins reveal their differential requirements for compensating age-dependent alterations in motor behavior and dopamine metabolism. Neurobiol. Aging 46, 107–112. doi:10.1016/j.neurobiolaging.2016.06.020

Devine, M.J., Ryten, M., Vodicka, P., Thomson, A.J., Burdon, T., Houlden, H., Cavaleri, F., Nagano, M., Drummond, N.J., Taanman, J.-W., Schapira, A.H., Gwinn, K., Hardy, J., Lewis, P.A., Kunath, T., 2011. Parkinson’s disease induced pluripotent stem cells with triplication of the α-synuclein locus. Nat. Commun. 2, 440. doi:10.1038/ncomms1453

Emani, M.R., Närvä, E., Stubb, A., Chakroborty, D., Viitala, M., Rokka, A., Rahkonen, N., Moulder, R., Denessiouk, K., Trokovic, R., Lund, R., Elo, L.L., Lahesmaa, R., 2015. The L1TD1 protein interactome reveals the importance of post-transcriptional regulation in human pluripotency. Stem Cell Rep. 4, 519–528. doi:10.1016/j.stemcr.2015.01.014

Enriquez-Barreto, L., Palazzetti, C., Brennaman, L.H., Maness, P.F., Fairén, A., 2012. Neural cell adhesion molecule, NCAM, regulates thalamocortical axon pathfinding and the organization of the cortical somatosensory representation in mouse. Front. Mol. Neurosci. 5, 76. doi:10.3389/fnmol.2012.00076

Flierl, A., Oliveira, L.M.A., Falomir-Lockhart, L.J., Mak, S.K., Hesley, J., Soldner, F., Arndt-Jovin, D.J., Jaenisch, R., Langston, J.W., Jovin, T.M., Schüle, B., 2014. Higher vulnerability and stress sensitivity of neuronal precursor cells carrying an alpha-synuclein gene triplication. PLoS One 9, e112413. doi:10.1371/journal.pone.0112413

Gleeson, J.G., Lin, P.T., Flanagan, L.A., Walsh, C.A., 1999. Doublecortin is a microtubule-associated protein and is expressed widely by migrating neurons. Neuron 23, 257–271. doi:10.1016/s0896-6273(00)80778-3

Greten-Harrison, B., Polydoro, M., Morimoto-Tomita, M., Diao, L., Williams, A.M., Nie, E.H., Makani, S., Tian, N., Castillo, P.E., Buchman, V.L., Chandra, S.S., 2010. αβγ-Synuclein triple knockout mice reveal age-dependent neuronal dysfunction. Proc. Natl. Acad. Sci. USA 107, 19573–19578. doi:10.1073/pnas.1005005107

Hardy, R.J., Loushin, C.L., Friedrich, V.L., Chen, Q., Ebersole, T.A., Lazzarini, R.A., Artzt, K., 1996. Neural cell type-specific expression of QKI proteins is altered in quakingviable mutant mice. J. Neurosci. 16, 7941–7949.

Hitoshi, N., Ken-ichi, Y., Jun-ichi, M., 1991. Efficient selection for high-expression transfectants with a novel eukaryotic vector. Gene 108, 193–199. doi:10.1016/0378-1119(91)90434-D

Hu, N., Strobl-Mazzulla, P., Sauka-Spengler, T., Bronner, M.E., 2012. DNA methyltransferase3A as a molecular switch mediating the neural tube-to-neural crest fate transition. Genes Dev. 26, 2380– 2385. doi:10.1101/gad.198747.112

Hussein, S.M., Batada, N.N., Vuoristo, S., Ching, R.W., Autio, R., Närvä, E., Ng, S., Sourour, M., Hämäläinen, R., Olsson, C., Lundin, K., Mikkola, M., Trokovic, R., Peitz, M., Brüstle, O., Bazett-Jones, D.P., Alitalo, K., Lahesmaa, R., Nagy, A., Otonkoski, T., 2011. Copy number variation and selection during reprogramming to pluripotency. Nature 471, 58–62. doi:10.1038/nature09871

Ichtchenko, K., Hata, Y., Nguyen, T., Ullrich, B., Missler, M., Moomaw, C., Südhof, T.C., 1995. Neuroligin 1: a splice site-specific ligand for beta-neurexins. Cell 81, 435–443. doi:10.1016/0092-8674(95)90396-8

Izant, J.G., McIntosh, J.R., 1980. Microtubule-associated proteins: a monoclonal antibody to MAP2 binds to differentiated neurons. Proc. Natl. Acad. Sci. USA 77, 4741–4745. doi:10.1073/pnas.77.8.4741

Kennedy, T.E., Serafini, T., de la Torre, J.R., Tessier-Lavigne, M., 1994. Netrins are diffusible chemotropic factors for commissural axons in the embryonic spinal cord. Cell 78, 425–435. doi:10.1016/0092-8674(94)90421-9

Liew, C.-G., Draper, J.S., Walsh, J., Moore, H., Andrews, P.W., 2007. Transient and stable transgene expression in human embryonic stem cells. Stem Cells 25, 1521–1528. doi:10.1634/stemcells.2006-0634

Lin, L., Göke, J., Cukuroglu, E., Dranias, M.R., VanDongen, A.M.J., Stanton, L.W., 2016. Molecular Features Underlying Neurodegeneration Identified through In Vitro Modeling of Genetically Diverse Parkinson’s Disease Patients. Cell Rep. 15, 2411–2426. doi:10.1016/j.celrep.2016.05.022

Liu, J., Jones, K.L., Sumer, H., Verma, P.J., 2009. Stable transgene expression in human embryonic stem cells after simple chemical transfection. Mol. Reprod. Dev. 76, 580–586. doi:10.1002/mrd.20983

Love, M.I., Huber, W., Anders, S., 2014. Moderated estimation of fold change and dispersion for RNA-seq data with DESeq2. Genome Biol. 15, 550–550. doi:10.1186/s13059-014-0550-8

Madan, B., Madan, V., Weber, O., Tropel, P., Blum, C., Kieffer, E., Viville, S., Fehling, H.J., 2009. The pluripotency-associated gene Dppa4 is dispensable for embryonic stem cell identity and germ cell development but essential for embryogenesis. Mol. Cell. Biol. 29, 3186–3203. doi:10.1128/MCB.01970-08

Mariani, J., Simonini, M.V., Palejev, D., Tomasini, L., Coppola, G., Szekely, A.M., Horvath, T.L., Vaccarino, F.M., 2012. Modeling human cortical development in vitro using induced pluripotent stem cells. Proc. Natl. Acad. Sci. USA 109, 12770–12775. doi:10.1073/pnas.1202944109

Mayshar, Y., Ben-David, U., Lavon, N., Biancotti, J.-C., Yakir, B., Clark, A.T., Plath, K., Lowry, W.E., Benvenisty, N., 2010. Identification and classification of chromosomal aberrations in human induced pluripotent stem cells. Cell Stem Cell 7, 521–531. doi:10.1016/j.stem.2010.07.017

Nichols, J., Zevnik, B., Anastassiadis, K., Niwa, H., Klewe-Nebenius, D., Chambers, I., Schöler, H., Smith, A., 1998. Formation of pluripotent stem cells in the mammalian embryo depends on the POU transcription factor Oct4. Cell 95, 379–391. doi:10.1016/s0092-8674(00)81769-9

Oliveira, L.M.A., Falomir-Lockhart, L.J., Botelho, M.G., Lin, K.H., Wales, P., Koch, J.C., Gerhardt, E., Taschenberger, H., Outeiro, T.F., Lingor, P., Schüle, B., Arndt-Jovin, D.J., Jovin, T.M., 2015. Elevated α-synuclein caused by SNCA gene triplication impairs neuronal differentiation and maturation in Parkinson’s patient-derived induced pluripotent stem cells. Cell Death Dis. 6, e1994. doi:10.1038/cddis.2015.318

Oyler, G.A., Higgins, G.A., Hart, R.A., Battenberg, E., Billingsley, M., Bloom, F.E., Wilson, M.C., 1989. The identification of a novel synaptosomal-associated protein, SNAP-25, differentially expressed by neuronal subpopulations. J. Cell Biol. 109, 3039–3052. doi:10.1083/jcb.109.6.3039

Polymeropoulos, M.H., Lavedan, C., Leroy, E., Ide, S.E., Dehejia, A., Dutra, A., Pike, B., Root, H., Rubenstein, J., Boyer, R., Stenroos, E.S., Chandrasekharappa, S., Athanassiadou, A., Papapetropoulos, T., Johnson, W.G., Lazzarini, A.M., Duvoisin, R.C., Di Iorio, G., Golbe, L.I., Nussbaum, R.L., 1997. Mutation in the alpha-synuclein gene identified in families with Parkinson’s disease. Science 276, 2045–2047. doi:10.1126/science.276.5321.2045

Prots, I., Grosch, J., Brazdis, R.-M., Simmnacher, K., Veber, V., Havlicek, S., Hannappel, C., Krach, F., Krumbiegel, M., Schütz, O., Reis, A., Wrasidlo, W., Galasko, D.R., Groemer, T.W., Masliah, E., Schlötzer-Schrehardt, U., Xiang, W., Winkler, J., Winner, B., 2018. α-Synuclein oligomers induce early axonal dysfunction in human iPSC-based models of synucleinopathies. Proc. Natl. Acad. Sci. USA 115, 7813–7818. doi:10.1073/pnas.1713129115

Raciti, M., Granzotto, M., Duc, M.D., Fimiani, C., Cellot, G., Cherubini, E., Mallamaci, A., 2013. Reprogramming fibroblasts to neural-precursor-like cells by structured overexpression of pallial patterning genes. Mol. Cell. Neurosci. 57, 42–53. doi:10.1016/j.mcn.2013.10.004

Radine, C., Peters, D., Reese, A., Neuwahl, J., Budach, W., Jänicke, R.U., Sohn, D., 2020. The RNA-binding protein RBM47 is a novel regulator of cell fate decisions by transcriptionally controlling the p53-p21-axis. Cell Death Differ. 27, 1274–1285. doi:10.1038/s41418-019-0414-6

Raudvere, U., Kolberg, L., Kuzmin, I., Arak, T., Adler, P., Peterson, H., Vilo, J., 2019. g:Profiler: a web server for functional enrichment analysis and conversions of gene lists (2019 update). Nucleic Acids Res. 47, W191–W198. doi:10.1093/nar/gkz369

Ross, O.A., Braithwaite, A.T., Skipper, L.M., Kachergus, J., Hulihan, M.M., Middleton, F.A., Nishioka, K., Fuchs, J., Gasser, T., Maraganore, D.M., Adler, C.H., Larvor, L., Chartier-Harlin, M.-C., Nilsson, C., Langston, J.W., Gwinn, K., Hattori, N., Farrer, M.J., 2008. Genomic investigation of alpha-synuclein multiplication and parkinsonism. Ann. Neurol. 63, 743–750. doi:10.1002/ana.21380

Schneider, B.L., Seehus, C.R., Capowski, E.E., Aebischer, P., Zhang, S.-C., Svendsen, C.N., 2007. Over-expression of alpha-synuclein in human neural progenitors leads to specific changes in fate and differentiation. Hum. Mol. Genet. 16, 651–666. doi:10.1093/hmg/ddm008

Shi, Y., Kirwan, P., Livesey, F.J., 2012. Directed differentiation of human pluripotent stem cells to cerebral cortex neurons and neural networks. Nat. Protoc. 7, 1836–1846. doi:10.1038/nprot.2012.116

Singh Dolt, K., Hammachi, F., Kunath, T., 2017. Modeling Parkinson’s disease with induced pluripotent stem cells harboring α-synuclein mutations. Brain Pathol. 27, 545–551. doi:10.1111/bpa.12526

Singleton, A.B., Farrer, M., Johnson, J., Singleton, A., Hague, S., Kachergus, J., Hulihan, M., Peuralinna, T., Dutra, A., Nussbaum, R., Lincoln, S., Crawley, A., Hanson, M., Maraganore, D., Adler, C., Cookson, M.R., Muenter, M., Baptista, M., Miller, D., Blancato, J., Hardy, J., Gwinn-Hardy, K., 2003. alpha-Synuclein locus triplication causes Parkinson’s disease. Science 302, 841. doi:10.1126/science.1090278

Specht, C.G., Schoepfer, R., 2001. Deletion of the alpha-synuclein locus in a subpopulation of C57BL/6J inbred mice. BMC Neurosci. 2, 11. doi:10.1186/1471-2202-2-11

Spillantini, M.G., Schmidt, M.L., Lee, V.M., Trojanowski, J.Q., Jakes, R., Goedert, M., 1997. Alpha-synuclein in Lewy bodies. Nature 388, 839–840. doi:10.1038/42166

Stagi, M., Fogel, A.I., Biederer, T., 2010. SynCAM 1 participates in axo-dendritic contact assembly and shapes neuronal growth cones. Proc. Natl. Acad. Sci. USA 107, 7568–7573. doi:10.1073/pnas.0911798107

Strehl, S., Glatt, K., Liu, Q.M., Glatt, H., Lalande, M., 1998. Characterization of two novel protocadherins (PCDH8 and PCDH9) localized on human chromosome 13 and mouse chromosome 14. Genomics 53, 81–89. doi:10.1006/geno.1998.5467

Vallot, C., Huret, C., Lesecque, Y., Resch, A., Oudrhiri, N., Bennaceur-Griscelli, A., Duret, L., Rougeulle, C., 2013. XACT, a long noncoding transcript coating the active X chromosome in human pluripotent cells. Nat. Genet. 45, 239–241. doi:10.1038/ng.2530

Vierbuchen, T., Ostermeier, A., Pang, Z.P., Kokubu, Y., Südhof, T.C., Wernig, M., 2010. Direct conversion of fibroblasts to functional neurons by defined factors. Nature 463, 1035–1041. doi:10.1038/nature08797

Wiedenmann, B., Franke, W.W., 1985. Identification and localization of synaptophysin, an integral membrane glycoprotein of Mr 38,000 characteristic of presynaptic vesicles. Cell 41, 1017–1028. doi:10.1016/s0092-8674(85)80082-9

Winner, B., Regensburger, M., Schreglmann, S., Boyer, L., Prots, I., Rockenstein, E., Mante, M., Zhao, C., Winkler, J., Masliah, E., Gage, F.H., 2012. Role of α-synuclein in adult neurogenesis and neuronal maturation in the dentate gyrus. J. Neurosci. 32, 16906–16916. doi:10.1523/JNEUROSCI.2723-12.2012

Wunderle, V.M., Critcher, R., Ashworth, A., Goodfellow, P.N., 1996. Cloning and characterization of SOX5, a new member of the human SOX gene family. Genomics 36, 354–358. doi:10.1006/geno.1996.0474

Zasso, J., Ahmed, M., Cutarelli, A., Conti, L., 2018. Inducible Alpha-Synuclein Expression Affects Human Neural Stem Cells’ Behavior. Stem Cells Dev. 27, 985–994. doi:10.1089/scd.2018.0011

