## Supplementary data for "An isogenic collection of pluripotent stem cell lines with elevated α-synuclein expression"

### Cortical neuronal differentiation is not impaired by elevated levels of α-synuclein in human pluripotent stem cells

#### Supplementary Figure 1

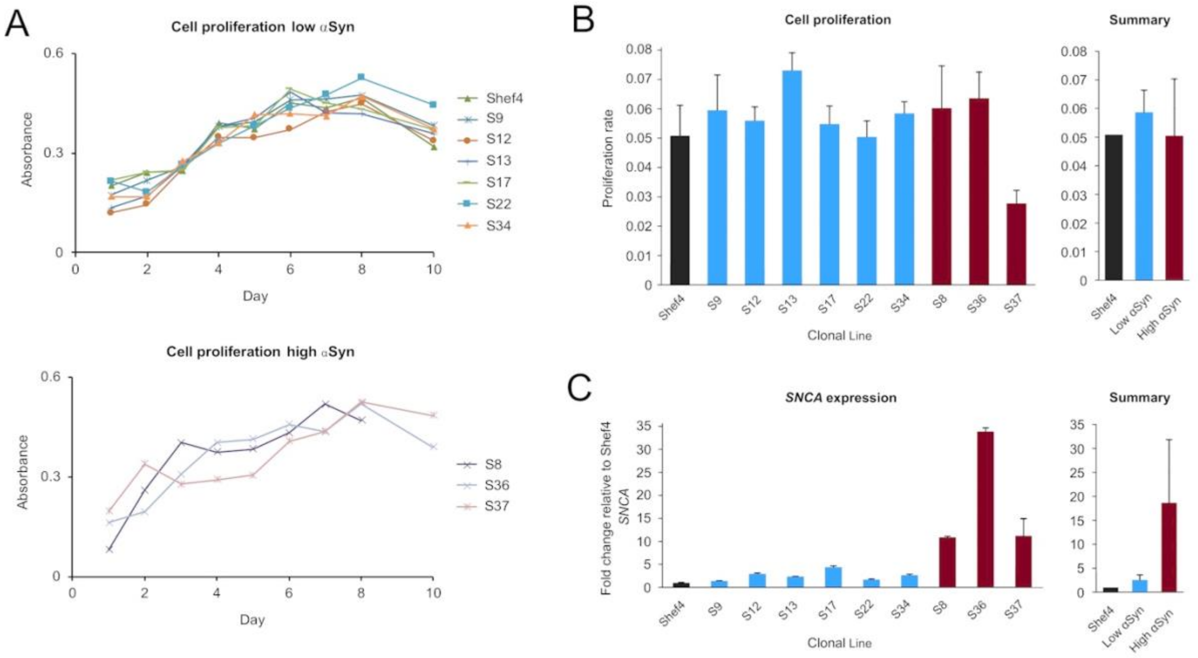


**Supplementary Figure 1. Proliferation rate of low and high αSyn hESC lines** (A) Proliferation curves representing cell proliferation, as measured at daily time points over the course of 10 days using the MTS assay for low αSyn cell lines (top) and high αSyn cell lines (bottom). A single sample per line was tested in three technical repeats and mean absorbance for each cell line was at 490 nm was plotted. n = 3 for each cell line (N. B. Data for S8 only obtained over an 8 day period). (B) Graph of proliferation rate of individual cell lines (left), and as an average of each group (right). Rate is calculated based on the slope of the linear trend line of the data obtained between Day 1 and Day 6 from the curves in (A). Data is shown for the parental line Shef4 (black), low *SNCA* expression cell lines (blue), and high *SNCA* expression lines (red). (C) Graph of *SNCA* expression level of individual cell lines shown as a fold-change relative to expression in the parental hESC Shef4 line (left), and as an average of each group (right). Error bars represent the SEM.
